# Fitting penalized regressions on very large genetic data using snpnet and bigstatsr

**DOI:** 10.1101/2020.10.30.362079

**Authors:** Florian Privé, Bjarni J. Vilhjálmsson, Hugues Aschard

**Author notes:** To whom correspondence should be addressed., Contact: •, •.

## Abstract

Both R packages snpnet and bigstatsr allow for fitting penalized regressions on individual-level genetic data as large as the UK Biobank. Here we benchmark bigstatsr against snpnet for fitting penalized regressions on large genetic data. We find bigstatsr to be an order of magnitude faster than snpnet when applied to the UK Biobank data (from 4.5x to 35x). We also discuss the similarities and differences between the two packages, provide theoretical insights, and make recommendations on how to fit penalized regressions in the context of genetic data.

## Introduction

Penalized regression with L1 penalty, also known as “lasso”, has been widely used since it proved to be an effective method for simultaneously performing variable selection and model fitting (Tibshirani 1996). R package glmnet is a popular software to fit the lasso efficiently (Friedman *et al.* 2010). However, glmnet cannot handle very large datasets such as biobank-scale data that are now available in human genetics, where both the sample size and the number of variables are very large. One strategy used to run penalized regressions on such large datasets such as the UK Biobank (Bycroft *et al.* 2018) has been to apply a variable pre-selection step before fitting the lasso (Lello *et al.* 2018). Recently, authors of the glmnet package have developed a new R package, snpnet, to fit penalized regressions on the UK Biobank without having to perform any pre-filtering (Qian *et al.* 2020). Earlier, we developed two R packages for efficiently analyzing large-scale (genetic) data, namely bigstatsr and bigsnpr (Privé *et al.* 2018). We then specifically derived a highly efficient implementation of penalized linear and logistic regressions in R package bigstatsr, and showed how these functions were useful for genetic prediction with some applications to the UK Biobank (Privé *et al.* 2019). Here we would like to come back to some statements made in (Qian *et al.* 2020) and benchmark bigstatsr against snpnet for fitting penalized regressions on large genetic data. We re-investigate the similarities and differences between the penalized regression implementations of packages snpnet and bigstatsr. Through some theoretical expectations and empirical comparisons, we show that package bigstatsr is generally much faster than snpnet, as opposed to what is presented in Qian *et al.* (2020) where they used a small synthetic dataset. We also make more recommendations on how to fit penalized regressions in the context of genetic data.

## Methods

### Main motivation for snpnet

Before we can present the main motivation behind snpnet developed by Qian *et al.* (2020), let us recall how the lasso regression is fitted. Fitting the lasso consists in finding regression coefficients *β* that minimize the following regularized loss function

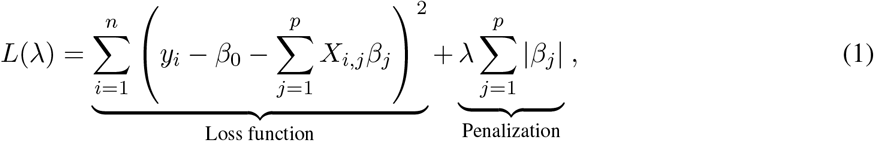

where *X* denotes the matrix composed of *p* (standardized) genotypes and possible covariates (e.g. sex, age and principal components) for *n* individuals, *y* is the (continuous) trait to predict, *λ* (> 0) is a regularization hyper-parameter that control the strength of the penalty. For a sequence of *λ*, one can find argmin_*β*_ *L*(*λ*) using cyclical coordinate descent (Friedman *et al.* 2010). To speed up the coordinate descent, one can use sequential strong rules for discarding lots of variables, i.e. setting lots of *β*_*j*_ to 0, a priori (Tibshirani *et al.* 2012). Therefore the cyclical coordinate descent used to solve the lasso can be performed in a subset of the data only thanks to these strong rules. However, the main drawback of these strong rules is that they require checking Karush-Kuhn-Tucker (KKT) conditions a posteriori, usually in two phases. These KKT conditions are first checked in the ever-active set, i.e. the set of all variables *j* with *β*_*j*_ ≠ 0 for any previous *λ*. Then, the cyclical coordinate descent has to be rerun while adding the new variables that do not satisfy these KKT conditions (if any). In a second phase, the KKT conditions are also checked for all the remaining variables, i.e. the ones not in the ever-active set. This last step requires to pass over the whole dataset at least once again for every *λ* tested. Then, when the available random access memory (RAM) is not large enough to cache the whole dataset, data has to be read from disk, which can be extremely time consuming. To alleviate this particular issue, Qian *et al.* (2020) have developed a clever approach called batch screening iterative lasso (BASIL) to be able to check these KKT conditions on the whole dataset only after having fitted solutions for many *λ*, instead of performing this operation for each *λ*. Hence, for very large datasets, the BASIL procedure enables to fit the exact lasso solution faster than when checking the KKT conditions for all variables at each *λ*, as performed in e.g. R package biglasso (Zeng and Breheny 2017).

### A more pragmatic approach in bigstatsr

Since our package bigstatsr is not using this BASIL procedure, reading Qian *et al.* (2020) could give the impression that fitting penalized regression with R package bigstatsr should be slow for very large datasets. We do check the KKT conditions for variables in the ever-active set, i.e. for a (small) subset of all variables only; this first checking is therefore fast. However, KKT conditions almost always hold when *p* > *n* (Tibshirani *et al.* 2012), which is particularly the case for the remaining variables in the second phase of checking. Because of this, we decided in Privé *et al.* (2019) to skip this second checking when designing functions `big_spLinReg` and `big_spLogReg` for fitting penalized regression on very large datasets in R package bigstatsr. Thanks to this approximation, these two functions effectively access all variables only once at the very beginning to compute the statistics used by the strong rules, and then access a subset of variables only (the ever-active set). As we will show, this means that fitting penalized regressions using the approximation we proposed in Privé *et al.* (2019) is computationally more efficient than using the BASIL procedure proposed by Qian *et al.* (2020), and yet provides as accurate predictors. Moreover, as bigstatsr uses memory-mapping, data that resides on disk is accessed only once from disk to memory and then stays in memory while there is no need to free memory, contrary to what could be read from Qian *et al.* (2020). Only when the ever-active set becomes very large, e.g. for very polygenic traits, memory can become an issue, but this extreme case would become a problem for package snpnet as well. Please refer to the Discussion section of Privé *et al.* (2019) for more details on these matters. In summary, bigstatsr effectively performs only one pass on the whole dataset while snpnet performs many passes, even though the number of passes in snpnet is reduced thanks to the BASIL approach. Moreover, bigstatsr still uses a single pass even when performing CMSA (a variant of cross-validation (CV), see figure 1) internally, whereas performing CV with snpnet would multiply the number of passes to the data by the number of folds used in the CV.

**Figure 1:**
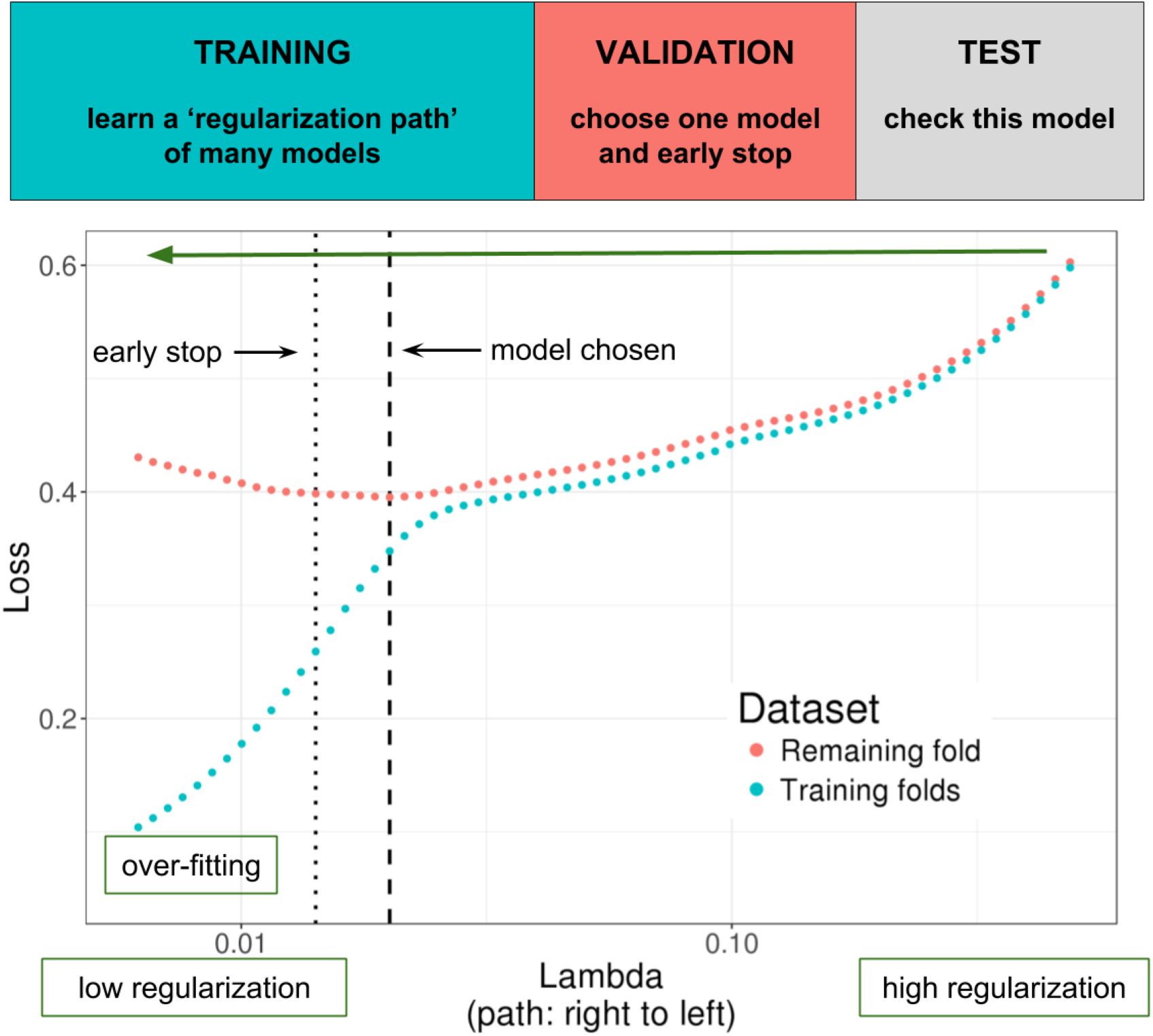
Illustration of one turn of the Cross-Model Selection and Averaging (CMSA) procedure. This figure comes from Privé *et al.* (2019); the Genetics Society of America has granted us permission to re-use it. First, this procedure separates the training set in *K* folds (e.g. 10 folds). Secondly, in turn, each fold is considered as an inner validation set (red) and the other (*K* − 1) folds form an inner training set (blue). A “regularization path” of models is trained on the inner training set and the corresponding predictions (scores) for the inner validation set are computed. The model that minimizes the loss on the inner validation set is selected. Finally, the *K* resulting models are averaged; this is different to standard cross-validation where the model is refitted on the whole training set using the best-performing hyper-parameters. We also use this procedure to derive an early stopping criterion so that the algorithm does not need to evaluate the whole regularization paths, making this procedure much faster.

### Data and methods for benchmark

As in Qian *et al.* (2020), we use the UK Biobank data (Bycroft *et al.* 2018), which is a large cohort of half a million individuals from the UK, for which we have access to both genotypes and multiple phenotypes (https://www.ukbiobank.ac.uk/). We apply some quality control filters to the genotyped data; we remove individuals with more than 10% missing values, variants with more than 1% missing values, variants having a minor allele frequency < 0.01, variants with P-value of the Hardy-Weinberg exact test < 10^−50^, and non-autosomal variants. We restrict individuals to the ones used for computing the principal components in the UK Biobank (Field 22020); these individuals are unrelated and have passed some quality control (Bycroft *et al.* 2018). We also restrict to the “White British” group defined by the UK Biobank (Field 22006) to get a set of genetically homogeneous individuals. These filters result in 337,475 individuals and 504,139 genotyped variants.

We use the same four phenotypes as used in Qian *et al.* (2020), namely height, body mass index (BMI), high cholesterol and asthma. We define height using field 50, BMI using field 21001, high cholesterol using field 20002 (“Non-cancer illness code, self-reported”). Asthma is defined using field 20002 as well as fields 40001, 40002, 41202 and 41204 (ICD10 codes); please see code for further details (Section “Software and code availability”). For height and BMI, L1-penalized linear regressions are fitted using function `big_spLinReg` from bigstatsr and using parameter `family=“gaussian”` in snpnet. For high cholesterol and asthma, L1-penalized logistic regressions are fitted using function `big_spLogReg` from bigstatsr and using parameter `family=“binomial”` in snpnet. We use sex (Field 22001), age (Field 21022), and the first 16 principal components (Field 22009) as unpenalized covariates when fitting the lasso models.

## Results

In the Methods section, we presented why we expect bigstatsr to be more efficient than snpnet. To practically support this claim, we perform comparisons for the four real traits used in the UK Biobank analyses of Qian *et al.* (2020). We compare the latest version of package snpnet (v0.3) with those of bigstatsr (v1.3) and bigsnpr (v1.5). We use similar quality controls as Qian *et al.* (2020) (Methods section). We also use the same splitting strategy: 20% test, 20% validation and 60% training. To use the same sets for bigstatsr as for snpnet, we use the same test set, use K=4 folds for training with bigstatsr while making sure one of them is composed of the same 20% of the data used for validation in snpnet. Moreover, we use penalty factors to effectively use unscaled genotypes in bigstatsr (see Discussion section), as performed by default in snpnet. This enables us to compare predictions from snpnet and bigstatsr using the exact same model and the same single validation fold. Note that, to make the most of the training set, bigstatsr uses CMSA (Figure 1) while Qian *et al.* (2020) propose to refit the model (on the whole training + validation) using the best *λ* identified using the validation set in snpnet. Also note that the parallelism used by snpnet and bigstatsr is different; snpnet relies on PLINK 2.0 to check KKT conditions in parallel, while bigstatsr parallelizes fitting of models from different folds and hyper-parameters. Because bigstatsr uses memory-mapping, the data is shared across processes and therefore it can fit these models in parallel without multiplying the memory needed. We allow for 16 cores to be used in these comparisons; bigstatsr effectively uses only 4 here (the number of folds). We allow for 128 GB of RAM for bigstatsr, but allow for 500 GB of RAM for snpnet because we had memory issues running it with only 128 GB.

Table 1 presents the results of this benchmark. Fitting lasso is 35 times faster using bigstatsr than using snpnet for high cholesterol, 29 times faster for asthma, 16 times faster for BMI, and 4.5 times faster for height. When using only one validation fold for choosing the best-performing *λ* and no refitting, snpnet and bigstatsr provide the same predictive performance, validating the use of the approximation in bigstatsr. When using the whole training set, i.e. when refitting in snpnet and using CMSA in bigstatsr, predictive performance is much higher than when the validation set is not used for training. For example, partial correlation for height is of 0.6116 with CMSA (i.e. using the average of 4 models) compared to 0.5856 when using only one of these models, showing how important it is to make the most of the training + validation sets. Also, CMSA can provide slightly higher predictive performance than the refitting strategy in snpnet, e.g. a partial correlation of 0.3324 vs 0.3221 for BMI.

**Table 1:** Benchmark of snpnet against bigstatsr in terms of predictive performance and computation time. Predictive performance is reported in terms of partial correlations between the polygenic scores and the phenotypes, residualized using the covariates. Timings are reported in minutes. Timings for snpnet report the training for 60% of the data (using the training set only) + the refitting for 80% of the data (using both the training and validation sets). Timings for bigstatsr report the time taken by the CMSA procedure (fitting K=4 models here).

| Trait | snpnet |  |  | bigstatsr |  |  |
| --- | --- | --- | --- | --- | --- | --- |
|  | Perf. (1 fold) | Perf. (refit) | Time | Perf. (1 fold) | Perf. (CMSA) | Time |
| Asthma | 0.1349 | 0.1438 | 188 + 101 | 0.1348 | 0.1493 | 10 |
| High cholesterol | 0.1254 | 0.1366 | 101 + 146 | 0.1257 | 0.1387 | 7 |
| BMI | 0.3031 | 0.3231 | 161 + 893 | 0.3018 | 0.3324 | 65 |
| Height | 0.5871 | 0.6106 | 409 + 715 | 0.5856 | 0.6116 | 249 |

## Discussion

We have found the BASIL approach derived in Qian *et al.* (2020) to be a clever approach that alleviates the I/O problem of other penalized regression implementations for very large datasets. BASIL makes significant and valuable contributions to the important problem of fitting penalized regression models efficiently. However, we also find that the implementation of BASIL in snpnet is still an order of magnitude slower than our package bigstatsr, which uses a simpler and more pragmatic approach (Privé *et al.* 2019). Hereinafter we also come back to some statements made in Qian *et al.* (2020) and provide more recommendations on how to best use penalized regression for deriving polygenic scores based on very large individual-level genetic data. This also enables us to highlight further similarities and differences between implementations from snpnet and bigstatsr.

First, in their UK Biobank applications, Qian *et al.* (2020) have tried using elastic-net regularization (a combination of L1 and L2 penalties) instead of lasso (only L1), i.e. introducing a new hyper-parameter *α* (0 < *α* < 1, with the special case of *α* = 1 being the L1 regularization). They show that L1 regularization is very effective for very large sample sizes, and elastic-net regularization is not needed in this case, which we have also experienced. Yet, in smaller sample sizes and for very polygenic architectures, we showed through extensive simulations that using lower values for *α* can significantly improve predictive performance (Privé *et al.* 2019). In Qian *et al.* (2020), they tried *α* ∈ {0.1, 0.5, 0.9, 1}; we recommend to use a grid on the log scale with smaller values (e.g. 1, 0.1, 0.01, 0.001, and even until 0.0001) for smaller sample sizes. Note that using a smaller *α* leads to a larger number of non-zero variables and therefore more time and memory required to fit the penalized regression. In functions `big_spLinReg` and `big_spLogReg` of R package bigstatsr, we allow to directly test many *α* values in parallel within the CMSA procedure. Therefore an optimal *α* value can be chosen automatically within the CMSA framework, without the need for more passes on the data.

Second, for large datasets, one should always use early-stopping. We have not found this to be emphasized enough in *Qian et al.* (2020). Indeed, while fitting the regularization path of decreasing *λ* values on the training set, one can monitor the predictive performance on the validation set, and stop early in the regularization path when the model starts to overfit (Figure 1). For large datasets, performance on the validation sets is usually very smooth and monotonic (before and after the minimum) along the regularization path, then one can safely stop very early, e.g. after the second iteration for which prediction becomes worse on the validation set. This corresponds to setting `n.abort=2` in bigstatsr and `stopping.lag=2` in snpnet. This is particularly useful because, when we move down the regularization path of *λ* values, more and more variables enter the model and the cyclical coordinate descent takes more and more time and memory. Therefore, the early-stopping criterion used in both bigstatsr and snpnet prevents from fitting very costly models, saving a lot of time and memory.

Third, Qian *et al.* (2020) recommend not to use scaled genotypes when applying lasso to genetic data. However, using scaled genotypes is common practice in genetics, and is the assumption behind models in popular software such as GCTA and LDSC (Yang *et al.* 2011; Bulik-Sullivan *et al.* 2015). Scaling genotypes assumes that, on average, all variants explain the same amount of variance and that low-frequency variants have larger effects. Speed *et al.* (2012) argued that this assumption might not be reasonable and proposed another model: 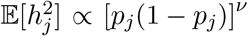, where 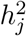 is the variance explained by variant *j* and *p*_*j*_ is its allele frequency. In Speed *et al.* (2017), they estimated *ν* to be between −0.25 and −0.5 for most traits. Note that scaling genotypes by dividing them by their standard deviations SD_*j*_ as done by default in bigstatsr assumes *ν* = −1 while not using any scaling as argued by Qian *et al.* (2020) assumes *ν* = 0. Therefore, using a trade-off between these two approaches can provide higher predictive performance and is therefore recommended (Zhang *et al.* 2020). In the case of L1 regularization, using a different scaling can be obtained by using different penalty factors *λ*_*j*_ in equation (1), which is an option available in both bigstatsr and snpnet. For example, using *λ*_*j*_ = 1/SD_*j*_ allows to effectively use unscaled genotypes. Recently, we have implemented a new parameter `power_scale` to allow for different scalings when fitting the lasso in bigstatsr. Note that a vector of values to try can be provided, and the best-performing scaling is automatically chosen within the CMSA procedure.

Fourth, Qian *et al.* (2020) write that bigstatsr “do not provide as much functionality as needed in [their] real-data application”, mainly because bigstatsr requires converting the input data and cannot handle missing values. It is true that bigstatsr uses an intermediate format, which is a simple on-disk matrix format accessed via memory-mapping. However, package bigsnpr provides fast parallel functions `snp_readBed2` for converting from ‘.bed’ files and `snp_readBGEN` for converting from imputed ‘.bgen’ files, the two formats used by the UK Biobank. For example, it took 6 minutes only to read from the UK biobank ‘.bed’ file used in this paper. We then used function `snp_fastImputeSimple` to impute by the variant means in 5 minutes only, which is also the imputation strategy used in snpnet. When reading imputed dosages instead, it takes less than one hour to access and convert 400K individuals over 1M variants using function `snp_readBGEN` with 15 cores, and less than three hours for 5M variants. When available, we recommend to directly read from ‘.bgen’ files to get dosages from external reference imputation. As for package snpnet, it uses the PLINK 2.0 ‘.pgen’ format, which is still under active development (in alpha testing, see https://www.cog-genomics.org/plink/2.0/formats#pgen). This format is not currently provided by the UK Biobank, and can therefore be considered as an intermediate format as well.

Fifth, Qian *et al.* (2020) point out the lack of flexibility of bigstatsr because it handles linear and logistic regressions but not Cox regression, and can only use standardized variables. Although Cox regression is not yet implemented in bigstatsr, it is one of the regression for which it is easy to derive strong rules (Tibshirani *et al.* 2012). As for scaling, we have seen how to effectively get unstandardized genotypes using different penalty factors, and we actually recommend to use an in-between scaling, or to test different scaling values within our CMSA framework using the new parameter `power_scale` in bigstatsr. Finally, when we developed R packages bigstatsr and bigsnpr for analyzing large scale genetic data, we separated functions in two packages because some functions are not specific to genetic data (the ones in bigstatsr). Therefore, when using our on-disk matrix format, one can store e.g. other omics data and has also access to a broad range of analysis tools provided by bigstatsr without any extra coding required, e.g. ultra-fast penalized regressions, association studies and principal component analysis (Privé *et al.* 2018).

## Software and code availability

All code used for this paper is available at https://github.com/privefl/paper2-PRS/tree/master/response-snpnet/code. R packages snpnet, bigstatsr and bigsnpr can be installed from GitHub. A tutorial on fitting penalized regressions with R package bigstatsr is available at https://privefl.github.io/bigstatsr/articles/penalized-regressions.html.

## Acknowledgements

This research has been conducted using the UK Biobank Resource under Application Number 41181. Authors would also like to thank GenomeDK and Aarhus University for providing computational resources and support that contributed to these research results.

